# Interleukin-11 promotes colonic epithelial repair after mechanical disruption

**DOI:** 10.64898/2026.05.29.727830

**Authors:** Tamami Suto, Takashi Nishina, Makoto Kashima, Yuta Suzuki, Souichirou Kubota, Yuji Goto, Shiro Yui, Hiroyasu Nakano, Katsuhide Okunishi

## Abstract

The intestinal epithelium relies on rapid repair to maintain homeostasis after injury, and dysregulation of this process contributes to the pathogenesis of inflammatory bowel disease and colorectal cancer. Interleukin-11 (IL-11), a fibroblast-derived cytokine elevated in these diseases, has well-documented effects on stromal cells, but its direct action on intestinal epithelial cells remains poorly characterized. Here, we used mouse colon organoids as an isolated epithelial system to directly examine the effects of IL-11 on epithelial cells. IL-11 stimulation activated the canonical JAK/STAT3 pathway, as evidenced by increased STAT3 phosphorylation and *Socs3* induction in a concentration-dependent manner. In a pipetting-based mechanical disruption model, IL-11 significantly increased the number of organoids recovered. Although mechanical disruption dominated the overall transcriptional landscape, RNA-seq analysis identified coordinated upregulation of STAT3 target genes and proliferation-related pathways specifically in response to IL-11. Pharmacological inhibition of STAT3 attenuated the IL-11-induced promotion of organoid recovery, indicating that STAT3 signaling mediates the epithelial response to IL-11 and maintains organoid size under basal conditions. Together, these findings demonstrate that IL-11 directly promotes intestinal epithelial repair after mechanical disruption through STAT3-dependent signaling, providing a mechanistic basis for its protective role in acute colonic injury.

## Introduction

The intestinal epithelium undergoes continuous self-renewal, and its homeostasis is maintained through strict regulation of epithelial cell proliferation, differentiation, and apoptosis [1–3]. Disruption of this balance results in abnormal cell proliferation and barrier dysfunction, which are central to the pathogenesis of inflammatory bowel disease (IBD) and colorectal cancer [1,4]. Maintaining intestinal homeostasis requires complex crosstalk among epithelial cells, immune cells, and stromal cells, and fibroblast-derived cytokines have recently emerged as key regulators of this process [5–7].

Intestinal stem cells (ISCs) reside at the crypt base and are maintained by their surrounding niche, including stromal fibroblasts that provide key signals such as Wnt ligands, R-spondins, and BMPs [7]. Intestinal organoids—three-dimensional epithelial structures grown from ISCs—reproduce key features of the native epithelium and provide a powerful system to examine how niche-derived factors directly influence epithelial cells [8]. By supplementing organoids cultured with defined factors (EGF, Noggin, and R-spondin1) with specific cytokines or physical stress, researchers can isolate the direct effects of individual signals on epithelial responses [8].

Interleukin-11 (IL-11) is a member of the IL-6 cytokine family that signals through IL-11Rα1 and gp130, activating JAK-STAT3, MAPK, and AKT pathways [9]. Although IL-11 expression is low in healthy intestinal tissue, it is elevated in IBD and colorectal cancer, where IL-11-producing fibroblasts accumulate in inflamed and tumor-adjacent stroma [10]. Recent studies have further implicated IL-11 in tissue fibrosis and aging across multiple organs [11,12]. In the intestine, IL-11 exhibits context-dependent dual roles: *Il11ra1*^-/-^ and *Il11*^-/-^ mice show exacerbated DSS-induced colitis but reduced AOM/DSS-induced tumor formation, suggesting protective effects during acute inflammation but pro-tumorigenic effects in chronic settings [10,13,14]. In addition, the evidence suggests that IL-11 may also act directly on epithelial cells: *Il11ra1* expression increases in *Apc*-mutant tumor organoids, which show enhanced STAT3/ERK phosphorylation upon IL-11 stimulation [10], and *Il11ra1*^*-/-*^ and *Il11*^*-/-*^ mice exhibit increased epithelial apoptosis during colitis [13]. Despite these observations, whether IL-11 directly regulates intestinal epithelial cell function under homeostatic or stress conditions remains poorly understood.

To address this gap, we used mouse colon organoids as an isolated epithelial system to examine the direct effects of IL-11 under defined culture conditions. Transcriptomic analysis revealed that IL-11 regulates epithelial gene expression via JAK-STAT3 signaling and that mechanical disruption-induced stress is required for robust transcriptional responses. These findings demonstrate that IL-11 directly modulates epithelial responses, providing a basis for understanding how IL-11 supports epithelial repair after injury.

## Materials and Methods

### Organoid Culture

Mouse colon epithelial organoids were established from C57BL/6 mice as described previously [10]. All experiments were performed according to the guidelines approved by the Institutional Animal Experiment Committee of Toho University School of Medicine (24-561). A detailed procedure was described in the supplementary methods.

### Organoid treatment

For pipetting-based mechanical fragmentation, organoids were released from Cultrex Basement Membrane Extract (BME), Type 2, PathClear (R&D Systems), mechanically disrupted, and reseeded. IL-11 (418-ML-025, R&D Systems) was added at the indicated concentrations. In addition, organoids were pre-treated with 10 μM C188-9 (S8605, Selleck) or vehicle (DMSO, final concentration 0.1%) for 60 min, followed by stimulation with 100 ng/mL IL-11 for 30 min. A detailed procedure was described in the supplementary methods.

### Western blotting

Western blotting was performed as previously described [10], and blots were quantified using Fiji software [15]. Antibodies and the detailed procedure are described in the Supplementary Methods.

### Image acquisition and analysis

Organoid images were acquired using an all-in-one microscope (BZ-X700, Keyence) and processed by maximum intensity projection. Organoid size and number were quantified using Fiji software [15]. The detailed procedure are described in the Supplementary Methods.

### RNA isolation and qPCR

Total RNAs were extracted from the indicated organoids by using RNA Plus (Takara), and cDNAs were synthesized with the RevertraAce qPCR RT Kit (Toyobo). qPCR was performed on a QuantStudio 3 system (Thermo Fisher Scientific) using *Hprt* as an internal control. Primer sequences are listed in the Supplementary Methods.

### RNA-seq and data analysis

RNA-seq libraries were prepared using the Lasy-Seq ver. 1.5 protocol [16], a 3’-directed sequencing method without mRNA enrichment, and sequenced on NovaSeq X Plus (Illumina). Reads were mapped to the mouse transcriptome (GRCm38) and quantified with Salmon. Differential expression was analyzed using TCC with FDR = 0.05. GSEA was performed using fgsea with gene sets from MSigDB, KEGG, and custom compilations. A detailed procedure was described in the Supplementary Methods. Raw and processed RNA-seq data have been deposited in the Gene Expression Omnibus (GEO) database under accession number GSE329523.

### Whole-mount staining of organoids and immunofluorescence

Whole-mount immunofluorescence staining, including signal amplification of pSTAT3, was performed as described in the Supplementary Methods. Images were obtained and analyzed using a BZ-X700 microscope and BZ-X Analyzer (Keyence).

### Statistical analysis

Statistical analyses were performed using R (version 4.5.1) and GraphPad Prism 11 (GraphPad Software). For organoid functional assays, one-way or two-way ANOVA with appropriate post-hoc tests was used as described in figure legends. For RNA-seq, differential expression was analyzed using the TCC package with Benjamini-Hochberg FDR correction (*q* < 0.05), and pathway enrichment was assessed by GSEA using fgseaMultilevel (eps = 0) with FDR correction. *p* < 0.05 or *p*adj < 0.05 was considered statistically significant. All data are presented as mean ± SD. Detailed statistical procedures are provided in the Supplementary Methods.

## Results

### IL-11 activates the JAK/STAT3 signaling pathway in established colonic organoids

To examine whether IL-11 activates the JAK/STAT3 signaling pathway in colonic epithelial cells, we treated established organoids with IL-11. Immunostaining (Figure 1A) and Western blot analysis (Figure 1B) revealed that IL-11 induced phosphorylation of STAT3 in a dose-dependent manner. Consistent with this, the expression of *Socs3*, a canonical STAT3 target gene, was upregulated by IL-11 in a concentration-dependent manner (Figure 1C). To assess the functional consequences of IL-11/STAT3 activation, organoids were treated with IL-11 for 2 days, and their size distribution was analyzed (Figure 1D, E). However, IL-11 did not significantly alter organoid number in either size category (500–5000 µm^2^ or >5000 µm^2^) compared with the control (Figure 1F). These results indicate that, despite robust activation of JAK/STAT3 signaling, IL-11 alone has a limited functional impact on established organoids under steady-state conditions, prompting us to investigate whether IL-11 functions in the context of epithelial injury.

**Figure 1.**
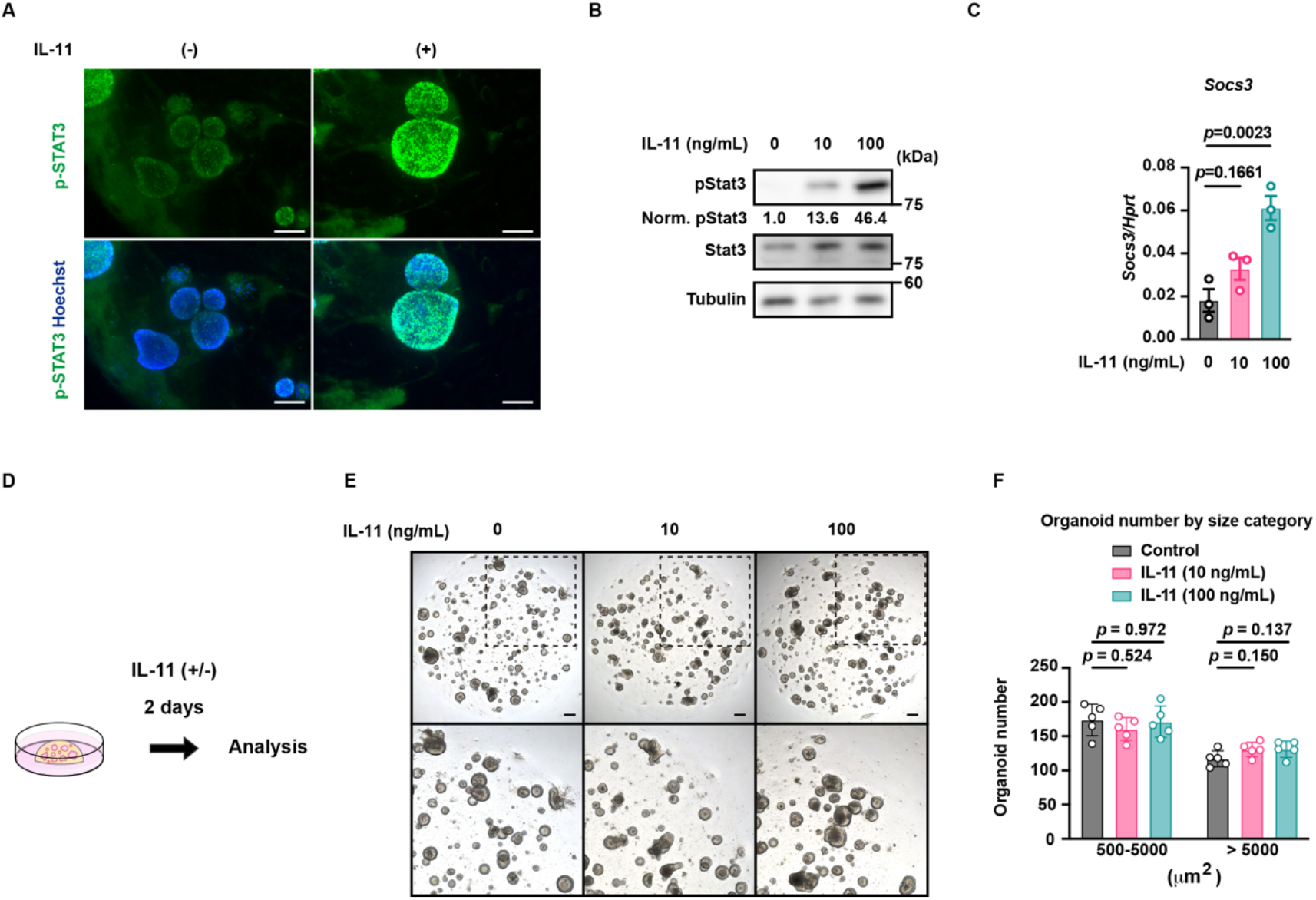
IL-11 activates the JAK/STAT3 signaling pathway in established colonic organoids. (A) Representative immunofluorescence images of organoids stained for phosphorylated STAT3 (p-STAT3, green) and Hoechst (blue) following IL-11 treatment (100 ng/mL, 30 min). Scale bars, 100 μm. Note that a low-level pSTAT3 signal is observed even in unstimulated organoids, indicating basal STAT3 activity. (B) Western blot analysis of p-STAT3, STAT3, and tubulin in organoids treated with IL-11 at the indicated concentrations. Numbers below the p-STAT3 blot indicate densitometric values normalized to total STAT3, relative to the untreated control. (C) *Socs3* mRNA expression in organoids treated with IL-11 at the indicated concentrations, measured by qPCR and normalized to *Hprt* (*n* = 3). Results are mean ± SD of technical triplicate samples. Statistical significance was determined by the one-way ANOVA with Dunnett’s multiple comparisons test. (D) Schematic of the experimental design for established organoid analysis. Established organoids were cultured with or without IL-11 (0, 10, or 100 ng/mL) for 2 days. (E) Representative brightfield images of organoids treated with IL-11 at the indicated concentrations for 2 days. Lower panels show magnified views of the boxed regions. Scale bars, 300 μm. (F) Number of organoids stratified by size category (500–5000 µm^2^ and >5000 µm^2^) after 2 days of culture. Symbols indicate individual wells; bars represent mean ± SD (*n* = 5 wells per condition). Statistical significance was determined by one-way ANOVA with Dunnett’s multiple-comparison test, performed within each size category. The data shown are representative of two independent experiments with similar results.

### IL-11 promotes organoid formation following *in vitro* mechanical disruption

Disruption of intercellular junctions and consequent loss of barrier integrity are central features of intestinal injury in inflammatory bowel disease [17], and such damage triggers regenerative reprogramming of the colonic epithelium *in vivo* [18]. Because pipetting similarly disrupts cell– cell contacts in organoids, we reasoned that this manipulation might recapitulate aspects of the *in vivo* injury response and used this model to examine whether IL-11 modulates this process. Organoids were dissociated by pipetting and treated with IL-11 for 3 days (Figure 2A). IL-11 concentration-dependently increased the total number of organoids (Figure 2B and C), with a more pronounced increase in small-sized (500–1000 μm^2^) organoids (Figure 2D). Indeed, the proportion of small-sized organoids was significantly increased by 100 ng/mL IL-11 (Figure 2E). These results indicate that IL-11 concentration-dependently promotes organoid formation after mechanical disruption; this effect was more pronounced in the small-sized population, despite a similar upward trend in the large-sized population.

**Figure 2.**
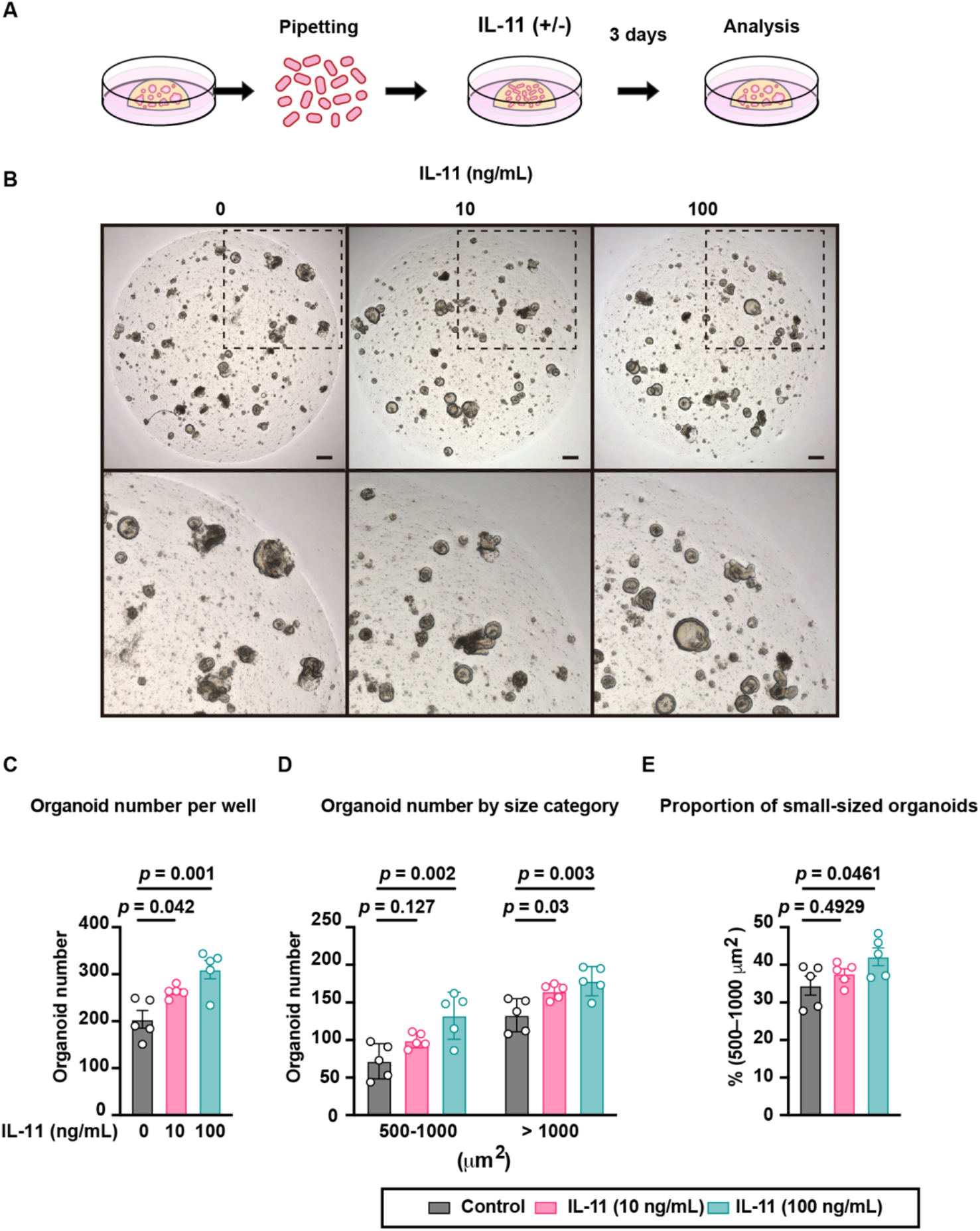
IL-11 promotes organoid formation following in vitro mechanical disruption. (A) Schematic of the experimental design. Organoids were dissociated by pipetting and cultured with or without IL-11 (0, 10, or 100 ng/mL) for 3 days. (B) Representative brightfield images at the indicated IL-11 concentrations. Lower panels show magnified views of the boxed regions. Scale bars, 300 μm. (C) Total organoid number per well after 3 days of culture. Symbols indicate individual wells; bars represent mean ± SD (*n* = 5 wells per condition). Statistical significance was determined by one-way ANOVA with Dunnett’s multiple comparisons test. (D) The same data shown in (C), stratified by size category (500–1000 µm^2^ and >1000 µm^2^). Symbols indicate individual wells; bars represent mean ± SD. Statistical significance was determined by one-way ANOVA with Dunnett’s multiple comparisons test, performed separately within each size category. (E) Proportion of small-sized organoids (500–1000 µm^2^) in each well, calculated as the percentage of small-sized organoids relative to the total organoid count per well. Symbols indicate individual wells; bars represent mean ± SD (*n* = 5 wells per condition). Statistical significance was determined by one-way ANOVA with Dunnett’s multiple comparisons test. The data shown are representative of two independent experiments with similar results.

### Mechanical disruption drives repair- and reprogramming-related transcriptional programs, while IL-11 selectively enhances a subset of proliferation- and STAT3-related genes

To characterize the transcriptional response to mechanical disruption and IL-11 treatment, organoids were disrupted by pipetting, treated with or without IL-11 for 3 hours, and subjected to RNA-seq analysis (Figure 3A). To assess whether this pipetting-based model recapitulates the epithelial injury response observed *in vivo*, we performed GSEA against mouse gene signatures of intestinal epithelial repair (mRepair) and fetal-like reprogramming (mFetal) reported during colitis [18]. Both the mRepair and mFetal signatures were significantly enriched at 3 hours post-pipetting (Figure 3B), indicating that mechanical disruption by pipetting activates transcriptional programs consistent with the *in vivo* epithelial repair and reprogramming response to injury. Principal component analysis (PCA) revealed that mechanical disruption was the dominant driver of transcriptional variance, with Control 0h samples clearly separated from the Pipetting groups along PC1 (14.6%). In contrast, pipetting samples clustered together regardless of IL-11 treatment, suggesting that IL-11 exerted only modest effects on the global transcriptional landscape (Figure 3C). Despite the subtle global transcriptional effect of IL-11, individual gene expression analysis revealed a subset of genes with suggestive differences (nominal *p* < 0.01) between IL-11-treated and untreated organoids (Figure 3D). To evaluate coordinated transcriptional shifts, we performed GSEA comparing IL-11-treated and IL-11-untreated organoids after mechanical disruption. GSEA revealed significant enrichment of cell proliferation-related genes, STAT3 target genes, and the intersection of proliferation and JAK-STAT pathway genes (Proliferation × JAK-STAT) in IL-11-treated organoids (Figure 3E; Supplementary Table S2 and S3). Heatmaps of leading-edge genes revealed distinct expression patterns between the IL-11-untreated and IL-11-treated groups (Figure 3F). These results indicate that IL-11 promotes epithelial proliferation and homeostasis by activating STAT3-dependent transcriptional programs.

**Figure 3.**
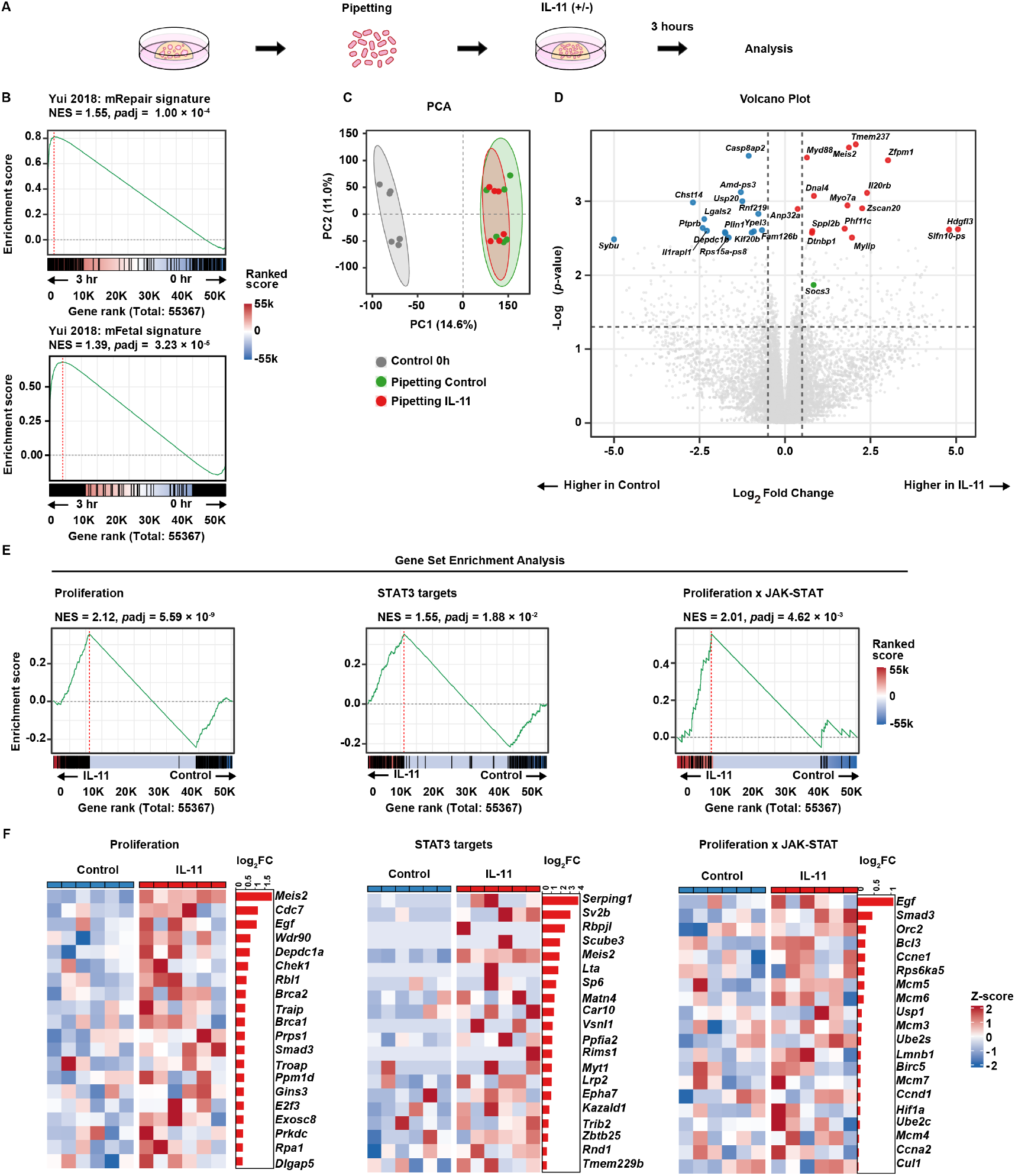
Mechanical disruption drives repair- and reprogramming-related transcriptional programs, while IL-11 selectively enhances a subset of proliferation- and STAT3-related genes. (A) Schematic of the experimental design. Colonic organoids were mechanically disrupted by pipetting and cultured for 3 hours with or without IL-11 (100 ng/mL). RNA-seq was performed on six samples per condition across two independent experiments. (B) GSEA enrichment plots for the mFetal (NES = 1.39, *p*adj = 3.23 × 10^−5^) and mRepair (NES = 1.55, *p*adj ≤ 1.00 × 10^−4^) signatures in IL-11-untreated organoids at 0 h vs. 3 h post-pipetting. Gene sets are described in Supplementary Methods. NES, normalized enrichment score. (C) PCA of transcriptional profiles from organoids untreated (0 h) or subjected to pipetting with or without IL-11 (100 ng/mL) (*n* = 6 per group). Each dot represents one biological sample; ellipses indicate 95% confidence regions. (D) Volcano plot of differentially expressed genes (pipetting with IL-11 vs. pipetting without IL-11, 3 h). Top 15 upregulated and downregulated genes (ranked by *p*-value, excluding predicted Gm genes) are labeled in red and blue, respectively. *Socs3* (green) is additionally labeled as a qPCR-validated STAT3 target. Dashed lines indicate *p* = 0.05 and log_2_FC = ±0.5. (E) GSEA enrichment plots for Proliferation, STAT3 targets, and Proliferation × JAK-STAT gene sets in IL-11-treated vs. control organoids after mechanical disruption. (F)Heatmaps of the top 20 leading-edge genes from each gene set in (E). Genes are ordered by log_2_FC (IL-11/Control). Colors represent row-wise Z-scores; bars indicate log_2_FC (red,upregulated; blue, downregulated in IL-11). Complete leading-edge gene lists are provided in Supplementary Table S3.

### The canonical IL-11/STAT3 signaling pathway promotes organoid formation after *in vitro* mechanical disruption

To examine whether IL-11 promotes organoid formation following mechanical disruption through STAT3 signaling, we first confirmed that the STAT3 inhibitor C188-9 [19] effectively blocked IL-11-induced STAT3 phosphorylation in colonic organoids (Figure 4A). We then dissociated organoids by pipetting and cultured them with vehicle or C188-9, in the absence or presence of IL-11, for 3 days (Figure 4B). The IL-11-induced increase in total organoid number was attenuated by C188-9 treatment, although this effect did not reach statistical significance (Figure 4C, D). Size-stratified analysis revealed that IL-11 significantly increased the number of small-sized organoids in the presence of DMSO, and this increase was abolished by C188-9 co-treatment (Figure 4E), indicating that STAT3 activity is required for IL-11-induced expansion of the small-sized population. In contrast, C188-9 significantly reduced the number of large-sized organoids both in the absence and presence of IL-11 (Figure 4E), suggesting that basal STAT3 signaling is required for the maintenance of larger organoids. Together, these results indicate that STAT3 activity contributes to both the IL-11-induced expansion of small-sized organoids and the maintenance of large-sized organoids.

**Figure 4.**
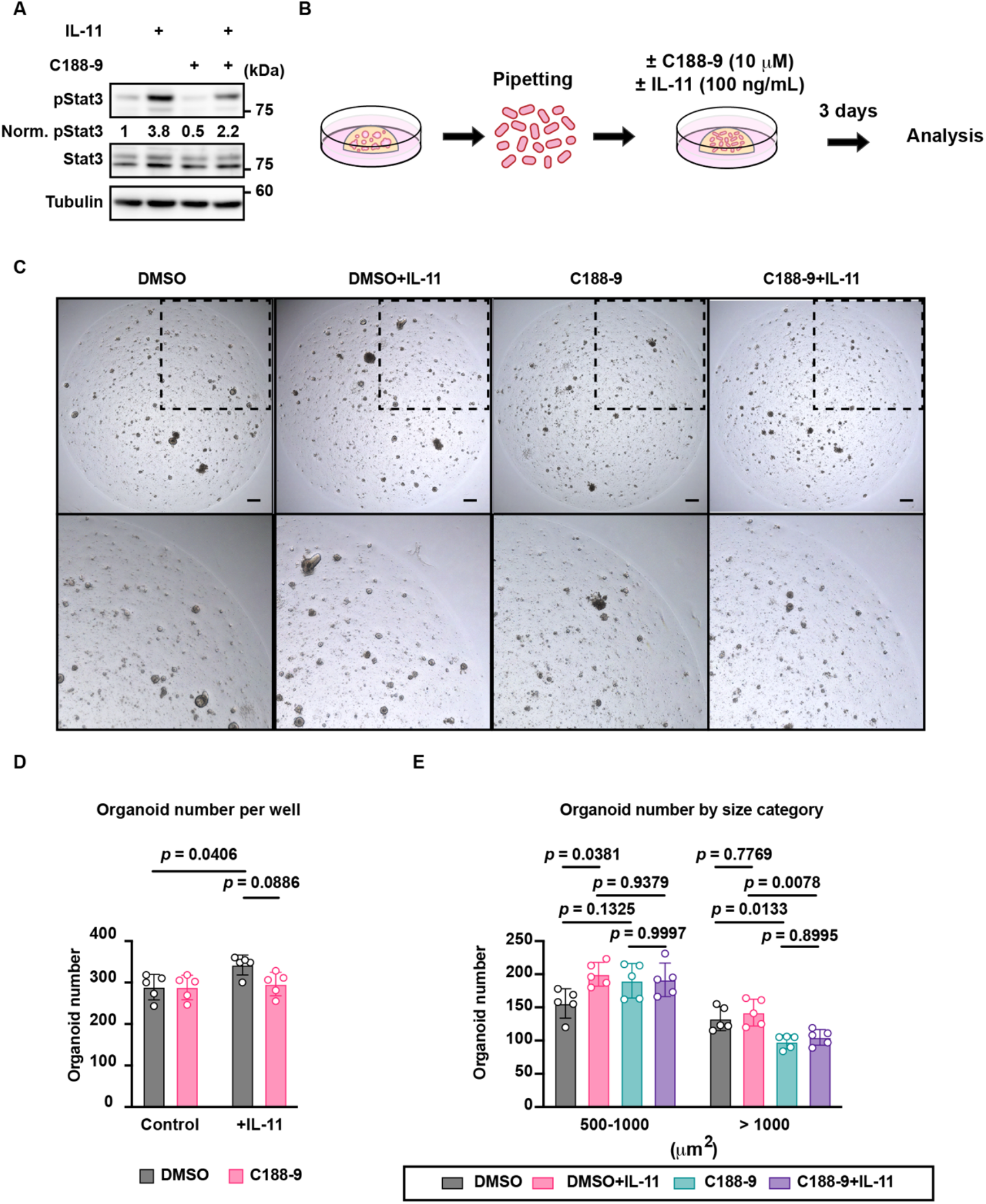
The canonical IL-11/STAT3 signaling pathway promotes organoid formation after *in vitro* mechanical disruption. (A) Western blot analysis of phosphorylated STAT3 (p-Stat3), total STAT3 (Stat3), and Tubulin in colonic organoids treated with vehicle (DMSO) or 10 μM C188-9, with or without 100 ng/mL IL-11 for 30 minutes. Numbers below the p-Stat3 blot indicate densitometric values normalized to total Stat3, relative to the DMSO control. (B) Schematic of the experimental procedure. Established colonic organoids were dissociated by pipetting and treated with vehicle (DMSO) or 10 μM C188-9, with or without 100 ng/mL IL-11. After 3 days of culture, organoids were analyzed for number and size distribution. (C) Representative brightfield images of organoids in each treatment condition after 3 days of culture. Lower panels show magnified views of the boxed regions. Scale bars, 300 μm. (D) Quantification of organoid number per well after pipetting and 3-day culture in the presence of vehicle (DMSO) or 10 μM C188-9, with or without 100 ng/mL IL-11. Bars represent mean ± SD; circles indicate individual replicates (*n* = 5 wells per condition). *p*-values were determined by two-way ANOVA followed by Tukey’s multiple comparisons test. (E) Organoid number stratified by size category (500–1000 µm^2^ and >1000 µm^2^) after pipetting and 3-day culture in the presence of vehicle (DMSO) or 10 μM C188-9, with or without 100 ng/mL IL-11. Symbols indicate individual wells; bars represent mean ± SD (*n* = 5 wells per condition). Statistical significance was determined by two-way ANOVA followed by Tukey’s multiple comparisons test, performed separately within each size category. The data shown are representative of two independent experiments with similar results.

## Discussion

Although IL-11 protects against DSS-induced colitis *in vivo* [13], whether it acts directly on intestinal epithelium through STAT3 signaling has remained unclear. In our *in vitro* model recapitulating the injury response, IL-11 treatment enhanced organoid recovery, as evidenced by increased organoid number, and the IL-11-induced increase in total organoid number was attenuated by STAT3 inhibition, and the maintenance of large-sized organoids was significantly impaired by C188-9 treatment. These results indicate that IL-11 promotes epithelial repair through canonical JAK/STAT3 signaling. Notably, C188-9 treatment significantly reduced the number of large-sized organoids independently of IL-11, suggesting that basal STAT3 activity, likely sustained by endogenous IL-6 family cytokines or other JAK/STAT activators in the culture environment, contributes to the maintenance of established organoids. This observation is consistent with the modest tendency of IL-11 alone to expand the large-sized organoid population under steady-state conditions, although this trend did not reach statistical significance.

Transcriptomic analysis revealed that mechanical disruption dominates the transcriptional landscape, while IL-11 induces coordinated regulation of injury-responsive pathways. Although no individual genes reached FDR significance, GSEA detected enrichment of STAT3 target, JAK-STAT, and proliferation-related gene sets at 3 hours post-injury, consistent with the known role of STAT3 in ISC-mediated regeneration [20,21]. These findings suggest that IL-11 promotes epithelial resilience through low-amplitude, coordinated regulation of STAT3 target genes, providing a mechanistic basis for its protective role in acute colitis [13]. Notably, pipetting-based dissociation alone activates regenerative transcriptional programs observed during *in vivo* repair, establishing it as a simple *in vitro* model for acute epithelial injury.

Several questions remain to be addressed. First, the molecular basis by which mechanical disruption confers IL-11 responsiveness is unclear; pipetting may modulate IL-11 receptor availability, signaling thresholds, or chromatin accessibility at STAT3 target loci. Second, the mechanisms underlying the size-dependent responses—STAT3-dependent expansion of small-sized organoids and STAT3-dependent maintenance of large-sized organoids—remain to be determined and likely reflect differences in cellular composition or proliferative state.

Addressing these questions will provide deeper insight into how epithelial cells integrate mechanical and cytokine signals during injury, with potential implications for IL-11-targeted therapies in IBD and colorectal cancer.

Our findings offer a mechanistic basis for IL-11’s paradoxical roles in intestinal disease. Oshima et al. demonstrated that STAT3 is indispensable for damage-induced crypt regeneration but dispensable for Wnt-driven intestinal tumorigenesis [22], suggesting that STAT3’s role is fundamentally context-dependent. Consistently, our data indicate that IL-11-mediated STAT3 activation in the acute injury setting promotes epithelial survival and regeneration, potentially limiting tissue damage in colitis [13]. In contrast, sustained IL-11 signaling in chronic inflammation or oncogenic settings may drive tumor progression [10,14,23]. Understanding this context-dependency may inform therapeutic strategies targeting IL-11 signaling in IBD and colorectal cancer.

## Data availability statement

The RNA-seq data generated in this study have been deposited in the NCBI Gene Expression Omnibus (GEO) database under accession number GSE329523 (https://www.ncbi.nlm.nih.gov/geo/query/acc.cgi?acc=GSE329523).

## Acknowledgements

This work has been supported in part by Grants-in-Aid for Scientific Research (B) 25K02536 (to TN); (B) 23K27398 (to HN); Grants-in-Aid for Scientific Research (C) 22K06932 (to TN), from the Japan Society for the Promotion of Science (JSPS); Extramural Collaborative Research Grant of Cancer Research Institute, Kanazawa University (to HN, and TN); and research grants from, the Takeda Science Foundation (to HN, MK, and TN), the Cell Research Foundation (to TN), the Daiichi Sankyo Foundation of Life Science (to TN), and the Mochida Foundation (to TN).

## CRediT authorship contribution statement

Tamami Suto: Investigation, Formal analysis, Visualization, Writing – original draft, Writing – review & editing.

Takashi Nishina: Conceptualization, Data curation, Investigation, Formal analysis, Funding acquisition, Project administration, Supervision, Visualization, Writing – original draft, Writing – review & editing.

Makoto Kashima: Funding acquisition, Methodology, Resources, Software, Writing – review & editing.

Yuta Suzuki: Investigation, Methodology, Writing – review & editing. Souichirou Kubota: Supervision, Writing – review & editing.

Yuji Goto: Supervision, Writing – review & editing. Shiro Yui: Data curation, Writing – review & editing.

Hiroyasu Nakano: Funding acquisition, Supervision, Writing – review & editing.

Katsuhide Okunishi: Supervision, Writing – review & editing.

## Declaration of generative AI and AI-assisted technologies in the manuscript preparation process

During the preparation of this work, the authors used Claude (Anthropic) to assist with English-language editing and improve the clarity of the manuscript. After using this tool, the authors reviewed and edited the content as needed and take full responsibility for the content of the published article.

