## Supplemental Methods for "Interleukin-11 promotes colonic epithelial repair after mechanical disruption"

**Supplementary Information**

**Author names**

Tamami Suto[1], Takashi Nishina[1]*, Makoto Kashima[2], Yuta Suzuki[2], Souichirou Kubota[3], Yuji Goto[3], Shiro Yui[4], Hiroyasu Nakano[5, 6], Katsuhide Okunishi[1]

[1]Department of Biochemistry, Faculty of Medicine, Toho University, Tokyo 143-8540, Japan

[2]Department of Biomolecular Science, Toho University, Chiba, 274-8510, Japan

[3]Department of Biology, Faculty of Science, Toho University, Chiba, 274-8510, Japan

[4]Center for Stem Cell and Regenerative Medicine, Institute of Biomedical Engineering, Institute of Science Tokyo, Tokyo 113-8510, Japan

[5]Unit of Host Defense, Faculty of Medicine, Toho University, Tokyo 143-8540, Japan

[6]Research Administration Organization, Toho University, Tokyo 143-8540, Japan

*Correspondence

**Corresponding Author**

Takashi Nishina, Ph.D.

Department of Biochemistry

Faculty of Medicine, Toho University

Tokyo 143-8540, Japan

**Supplementary methods**

**Organoid Culture**

Murine colonic organoids were embedded in Cultrex Basement Membrane Extract (BME), Type 2, PathClear (R&D Systems), and seeded into 48-well plates. The organoids were cultured in Advanced DMEM/F12 (Thermo Fisher Scientific) containing 10 mM HEPES (Thermo Fisher Scientific), 1 x GlutaMAX (Thermo Fisher Scientific), 1 x B27 (Thermo Fisher Scientific), 1 μM N-Acetylcysteine (NAC), murine epidermal growth factor (50 ng/mL, Thermo Fisher Scientific), murine Noggin (100 ng/mL, Peprotech), 0.5 mM A83-01 (Tocris), 3 μM SB202190 (Sigma), 1 μM nicotinamide (Sigma), Afamin and Wnt3A-condition medium (CM) (final concentration of 50%) which was provided by J. Takagi [1] and and R-spondin 1-CM (final concentration of 10%) (Trevigen). For the first two days after subculturing, the medium was supplemented with 10 μM Y-27632 (Nacalai Tesque) to inhibit anoikis. The culture medium was replaced every two days, and organoids were subcultured every 5-7 days. Two independent experiments were performed on different days using passaged organoids from the same source. To evaluate growth efficiency, organoids were cultured with or without IL-11 for 2-3 days. The total number of organoids per well was manually counted from light microscopy images using Fiji (ImageJ). RNA-seq libraries were prepared simultaneously from all wells, yielding six samples per condition (n = 6) across the two experiments.

**Pipetting Model and IL-11 Stimulation**

Organoids were recovered from the BME Type 2 matrix using Cell Recovery Solution (R&D Systems) with incubation at 4°C for 20 min. During recovery, the organoids were mechanically dissociated into small fragments using low-retention tips (Platinum Soft Tip, BM Bio). After washing with PBS, the organoid pellet was resuspended in fresh BME Type 2 and re-seeded as 10 μL droplets per well. Following gelation, the organoids were stimulated with 100 ng/mL IL-11 in 300 μL of culture medium for 3 days. Bright-field images were captured using a BZ-X700 microscope (Keyence).

**Western blotting**

Cells were lysed in RIPA buffer as previously described [10], and lysates were subjected to SDS-PAGE and transferred onto polyvinylidene difluoride membranes (Millipore). The membranes were developed with Immobilon Western Chemiluminescent HRP Substrate (Millipore) and analyzed using an Amersham Biosciences Imager 600 (GE Healthcare). Blots were quantified using Fiji software [15]. The antibodies used in this study and the detailed procedure were described in the supplementary methods.

**Immunostaining and Fluorescence Tyramide Signal Amplification**

Organoids were seeded on sterile glass coverslips in a 1:1 mixture of culture medium and BME Type 2. Prior to stimulation, the cells were starved for 1 h in Advanced DMEM/F12 containing only 10 mM HEPES and 1 x GlutaMAX, followed by treatment with 100 ng/mL IL-11 for 30 min. The samples were fixed with 4% formaldehyde (Nacalai Tesque) for 15 min at room temperature. To quench endogenous peroxidase activity, the organoids were incubated in chilled 3% H2O2 in methanol at -20°C for 10 min. After blocking with 5% normal donkey serum and 0.5% Triton X-100 in 2% BSA-PBS for 30 min, the samples were incubated overnight at 4°C with the primary antibody p-STAT3 antibody (9132, Cell Signaling Technology). Then, organoids were stained with ImmPRESS-VR Horse anti-Rabbit IgG Polymer kit (MP-6401, Vector Laboratories) and Hoechst 33258. Fluorescence signals for p-STAT3 were subsequently amplified using the Opal 520 Reagent Pack (FP1487001KT, AKOYA Biosciences) according to the manufacturer's instructions. Finally, the coverslips were mounted using VECTASHIELD PLUS Antifade Mounting Medium (Vector Laboratories), and fluorescence images were captured with a BZ-X700 microscope (Keyence).

**RNA isolation and qPCR**

Total RNAs were extracted from the indicated organoids by using RNA Plus (Takara), and cDNAs were synthesized with the RevertraAce qPCR RT Kit (Toyobo). qPCR was performed on a QuantStudio 3 system (Thermo Fisher Scientific) using the ΔCt method with *Hprt* as an internal control. The following primers were used in this study: murine *Socs3*, 5 - ATGGTCACCCACAGCAAGTTT - 3 and 5 - TCCAGTAGAATCCGCTCTCCT -3 ; murine *Hprt*, 5 - AACAAAGTCTGGCCTGTATCCAA -3 and 5 - GCAGTACAGCCCCAAAATGG -3.

**RNA-Seq library preparation and sequencing**

RNA-Seq library preparation was conducted according to the Lasy-Seq ver. 1.5 protocol [2]. Total RNAs were reverse transcribed using an RT primer with index and SuperScript IV reverse transcriptase (Thermo Fisher Scientific). Then, all RT mixtures of the samples were pooled and purified using an equal volume of KAPA HyperPure Beads (F. Hoffmann-La Roche, Ltd.) according to the manufacturer's instructions. Second-strand synthesis was conducted on the pooled samples using RNaseH (5 U/µL, Enzymatics) and DNA polymerase I (10 U/µL, Enzymatics). To avoid the carryover of large amounts of rRNA, the mixture was treated with RNase T1 (Thermo Fisher Scientific). Then, purification was conducted with a 0.8× volume of KAPA HyperPure Beads. Fragmentation, end-repair, and A-tailing were conducted using 5× WGS Fragmentation Mix (Enzymatics). The Adapter for Lasy-Seq was ligated using 5× Ligation Mix (Enzymatics), and the adapter-ligated DNA was purified with a 0.8× volume of KAPA HyperPure Beads. Then, only cDNA fragments containing polyA (3' parts of mRNA) were labeled with one cycle PCR using biotin-conjugated P7 primer ([BioON]CAAGCAGAAGACGGCATACGAGAT) and KAPA HiFi HotStart ReadyMix (F. Hoffmann-La Roche, Ltd.) on MiniAmp Thermal Cycler (Applied Biosystems). The biotin-labeled DNA was purified with a 0.8× volume of KAPA HyperPure Beads, followed by elution with 16 µL of 10 mM Tris-HCl (pH 7.6). The biotin-labeled DNA was captured with Dynabeads M-280 Streptavidin (Thermo Fisher Scientific) in 1M NaCl and 8 mM Tris-HCl (pH 7.6), followed by two cycles of library amplification. 9.5 µL of the amplified DNA was used for optimization of PCR cycles for library amplification by qPCR using EvaGreen, 20× in water (Biotium), and the AriaMx Real-Time PCR System (Agilent Technologies). Finally, the library was amplified using KAPA HiFi HotStart ReadyMix on MiniAmp Thermal Cycler. The amplified library was purified with an equal volume of KAPA HyperPure Beads. One microliter of the library was then used for electrophoresis using a Bioanalyzer 2100 with the Agilent High Sensitivity DNA kit (Agilent Technologies) to assess quality. Then, 150-bp paired-end reads were sequenced using the NovaSeq X Plus (Illumina).

**Western blotting**

Cells were lysed in RIPA buffer (50 mM Tris-HCl, pH 8.0, 150 mM NaCl, 1% Nonidet P-40, 0.5% deoxycholate, 0.1% SDS, 25 mM b-glycerophosphate, 1 mM sodium orthovanadate, 1 mM sodium fluoride, 1 mM PMSF, 1 mg/mL aprotinin, and 1 mg/mL leupeptin). After centrifugation, cell lysates were subjected to SDS–PAGE and transferred onto polyvinylidene difluoride membranes (Millipore). The membranes were developed with Immobilon Western Chemiluminescent HRP Substrate (Millipore) and analyzed using an Amersham Biosciences Imager 600 (GE Healthcare). Blots were quantified using Fiji software[3]. The following antibodies used in this study were obtained from the indicated sources: anti-phospho-STAT3 (9145, Cell Signaling Technology, 1:1000), anti-STAT3 (SC-482, Santa Cruz, 1:1000), anti-tubulin (T5168, Sigma-Aldrich, 1:50,000), Peroxidase AffiniPure Donkey Anti-Rabbit IgG (H+L) (711-035-152, Jackson Immuno Research Laboratories Inc.) and Peroxidase AffiniPure Donkey Anti-Mouse IgG (H+L) (715-035-151, Jackson Immuno Research Laboratories Inc.) antibodies.

**Mapping and gene expression quantification**

Gene expression quantification was performed using read 1 sequences, which contain the cDNA-derived sequences in the Lasy-Seq protocol. The reads were processed with fastp (version 0.23.4) [4] using the following parameters: --trim_poly_x -w 20 --adapter_sequence=AGATCGGAAGAGCACACGTCTGAACTCCAGTCA --adapter_sequence_r2=AGATCGGAAGAGCGTCGTGTAGGGAAAGAGTGT -l 31. The trimmed reads were then mapped to the Mus musculus reference sequences. GRCm38.cdna.all.fa, using BWA mem (version 0.7.17-r1188) [5] with the default parameters. The read count for each gene was calculated with salmon using -l IU, which specifies the library type (version v1.10.2) [6]. Then, using R (version 4.5.1) (R Core Team, 2017), the sums of read counts per gene were calculated. Raw and processed RNA-seq data have been deposited in the Gene Expression Omnibus (GEO) database under accession number GSE329523.

**Gene Set Preparation for Pathway Analysis**

Gene sets for pathway enrichment analysis were compiled from multiple curated databases and recent literature to comprehensively assess biological processes relevant to intestinal epithelial cell homeostasis and IL-11 signaling. Standard pathway gene sets were obtained from MSigDB (Molecular Signatures Database) via the msigdbr package (version 25.1.1), including MSigDB Hallmark gene sets, KEGG pathways, Reactome pathways, and Gene Ontology Biological Process (GO: BP) terms. Custom gene sets were assembled for key biological processes and signaling pathways as follows: Cell proliferation genes (*n*=325) were assembled from MSigDB Hallmark gene sets (HALLMARK_G2M_CHECKPOINT, HALLMARK_E2F_TARGETS), GO: BP terms (GO:0008283, cell proliferation), and KEGG cell cycle pathway (mmu04110), encompassing cell cycle regulators (*Ccnd1*, *Cdk4*, *Cdk6*), checkpoint proteins (*Cdkn1a*, *Cdkn1b*), and proliferation markers (*Mki67*, *Pcna*). JAK-STAT signaling genes (n=755) were compiled from MSigDB Hallmark (HALLMARK_IL6_JAK_STAT3_SIGNALING), KEGG (mmu04630, JAK-STAT signaling pathway), Reactome pathways (R-MMU-449147, Interleukin signaling; R-MMU-913531, Interferon signaling), and GO:BP terms (GO:0007259, receptor signaling via JAK-STAT; GO:0019221, cytokine-mediated signaling), encompassing JAK kinases (*Jak1*, *Jak2*, *Jak3*, *Tyk2*), STAT transcription factors (Stat1-6), cytokine receptors, and negative regulators (*Socs1-3*, *Pias1*, *Pias3*, *Ptpn11*). STAT3 target genes (*n*=168) were compiled from ChIP-seq databases (ChEA, ENCODE), primary literature, and the TRRUST transcription factor database, including direct STAT3 target genes validated by chromatin immunoprecipitation studies. A composite gene set, "Proliferation × JAKSTAT," was generated by intersecting the Cell proliferation and JAK-STAT signaling gene sets to identify genes involved in JAK-STAT-mediated proliferative responses. All gene symbols were converted to official Mouse Genome Informatics (MGI) nomenclature, cross-referenced with Ensembl Mouse Genome annotation (GRCm38.101), and validated for expression in our RNA-seq dataset (genes with total read counts > 0 across all samples). To assess regenerative reprogramming, we incorporated two specific gene signatures described by Yui et al. (2018) [7]: the mFetal-like and mRepair signatures. The mFetal-like signature, representing the transcriptional program of fetal intestinal spheroids, was obtained from the Supplementary Table of the original study. The mRepair signature (genes upregulated in DSS-injured Sca-1$^{+}$ intestinal epithelial cells; logFC > 1, FDR < 0.05) was provided by the authors and is also listed in Supplementary Table S1. The complete gene lists for the three custom gene sets used in GSEA (Cell Proliferation, STAT3 Targets, and Proliferation × JAK-STAT) are provided in Supplementary Table S2.

**Gene Set Enrichment Analysis**

Gene set enrichment analysis (GSEA) was performed using the fgseaMultilevel function from the fgsea package (version 1.34.0) [8]. For each comparison, genes were ranked by the M-value (log2 fold change estimate) calculated from TCC differential expression analysis. The ranked gene lists were tested against all compiled gene sets, including MSigDB collections and custom gene sets described above. Gene sets containing 5-500 genes were included in the analysis to balance statistical power and biological interpretability. Enrichment significance was assessed using the adaptive multilevel splitting algorithm (eps = 0), which provides accurate estimation of low p-values without the resolution limit imposed by standard permutation testing. p-values were adjusted for multiple testing using the Benjamini-Hochberg procedure, and pathways with adjusted *p*-value (*p*adj) < 0.05 were considered significantly enriched.

**RNA-seq Data Visualization**

Volcano plots, principal component analysis plots, and GSEA enrichment plots were generated using ggplot2 (version 3.5.2). Heatmaps were created using ComplexHeatmap (version 2.24.0) and pheatmap (version 1.0.12). Z-score normalization was applied to the heatmap for visualization using hierarchical clustering with Ward's method (ward.D2). Additional data manipulation and visualization were performed using dplyr (version 1.1.4), tidyr (version 1.3.1), and scales (version 1.4.0). All figures were exported in SVG format for publication-quality visualization.

[1] E. Mihara, H. Hirai, H. Yamamoto, K. Tamura-Kawakami, M. Matano, A. Kikuchi, T. Sato, J.

Takagi, Active and water-soluble form of lipidated Wnt protein is maintained by a serum glycoprotein afamin/α-albumin, Elife 5 (2016). https://doi.org/10.7554/eLife.11621.
